# Impacts of predators, habitat, recruitment, and disease on soft-shell clams *Mya arenaria* and stout razor clams *Tagelus plebeius* in Chesapeake Bay

**DOI:** 10.1101/224071

**Authors:** Cassandra N. Glaspie, Rochelle D. Seitz, Matthew B. Ogburn, Christopher F. Dungan, Anson H. Hines

## Abstract

Soft-shell clams, *Mya arenaria*, and razor clams, *Tagelus plebeius*, in Chesapeake Bay have declined since the 1970s, with severe declines since the 1990s. These declines are likely caused by multiple factors including warming, predation, habitat loss, recruitment limitation, disease, and harvesting. A bivalve survey in Chesapeake Bay examined influential factors on bivalve populations, focusing on predation (crabs, fish, and cownose rays), habitat (mud, sand, gravel, shell, or seagrass), environment (temperature, salinity, and dissolved oxygen), recruitment, and disease. *M. arenaria* and *T. plebeius* were found more often in habitats with complex physical structures (seagrass, shell) than any other habitat. Pulses in bivalve density associated with recruitment were attenuated through the summer and fall when predators are most active, indicating that predators likely influence temporal dynamics in these species. Presence of *Mya arenaria,* which is near the southern extent of its range in Chesapeake Bay, was negatively correlated with water temperature. Recruitment of *M. arenaria* in Rhode River, MD, declined between 1980 and 2016. Infection by the parasitic protist *Perkinsus* sp. was associated with stressful environmental conditions, bivalve size, and environmental preferences of *Perkinsus* sp, but was not associated with bivalve densities. It is likely that habitat loss, low recruitment, and predators are major factors keeping *T. plebeius* and *M. arenaria* at low densities in Chesapeake Bay. Persistence at low densities may be facilitated by habitat complexity (presence of physical structures), whereas further reductions in habitats such as seagrass and shell hash could result in local extinction of these important bivalve species.

## INTRODUCTION

Soft-shell clams, *Mya arenaria*, and razor clams, *Tagelus plebeius*, in Chesapeake Bay have declined since the 1970s, with severe declines since the 1990s. These declines are likely caused by multiple factors including warming, predation, habitat loss, recruitment limitation, disease, and harvesting. A bivalve survey in Chesapeake Bay examined influential factors on bivalve populations, focusing on predation (crabs, fish, and cownose rays), habitat (mud, sand, gravel, shell, or seagrass), environment (temperature, salinity, and dissolved oxygen), recruitment, and disease. *M. arenaria* and *T. plebeius* were found more often in habitats with complex physical structures (seagrass, shell) than any other habitat. Pulses in bivalve density associated with recruitment were attenuated through the summer and fall when predators are most active, indicating that predators likely influence temporal dynamics in these species. Presence of *Mya arenaria,* which is near the southern extent of its range in Chesapeake Bay, was negatively correlated with water temperature. Recruitment of *M. arenaria* in Rhode River, MD, declined between 1980 and 2016. Infection by the parasitic protist *Perkinsus* sp. was associated with stressful environmental conditions, bivalve size, and environmental preferences of *Perkinsus* sp, but was not associated with bivalve densities. It is likely that habitat loss, low recruitment, and predators are major factors keeping *T. plebeius* and*M. arenaria* at low densities in Chesapeake Bay. Persistence at low densities may be facilitated by habitat complexity (presence of physical structures), whereas further reductions in habitats such as seagrass and shell hash could result in local extinction of these important bivalve species.

The soft-shell clam *Mya arenaria* and the stout razor clam *Tagelus plebeius* are both large, deep-burrowing bivalves that are harvested in the Chesapeake Bay for human consumption and for bait (Dungan et al. 2002, Homer et al. 2011). *Mya arenaria,* in particular, supports a large commercial fishery in the U.S., and it accounted for 14% of domestic commercial bivalve dollar value in 2015 (NMFS 2016). In Chesapeake Bay, *M. arenaria* has supported a commercial hydraulic dredge fishery in Virginia and Maryland starting in the early 1950s with the invention of the hydraulic dredge. Commercial clammers also harvest *T. plebeius* for eel and crab bait (Dungan et al. 2002, Homer et al. 2011).

Historically, *M. arenaria* and *T. plebeius* served important roles as biomass dominants that contributed substantially to the food web and water quality of Chesapeake Bay (Abraham & Dillon 1986, Eggleston et al. 1992, Seitz et al. 2001). *Mya arenaria* and *T. plebeius* are key prey for numerous commercially and recreationally important fish and crab species (Eggleston et al. 1992, Seitz et al. 2005, Fisher 2010). These large-bodied filter-feeding clams likely played a large role in filtration of the water column when they were abundant; *M. arenaria* in the Baltic Sea can filter the entire water column in less than a day (Forster & Zettler 2004), and filtration by *M. arenaria* in the Gulf of Maine increased water clarity in some lagoons where their densities were high (Thiet et al. 2014). Non-oyster bivalves have recently gained attention for water filtration as an ecosystem service in Chesapeake Bay (Gedan et al. 2014), but the value of filtration by thin-shelled clams has not been assessed.

In Chesapeake Bay, *M. arenaria* has been in decline since the early 1970s, with more pronounced declines since the 1990s, and this species now exists in Chesapeake Bay at record low levels (Fig. 1). Declines during and after 1972 are attributed to impacts from Tropical Storm Agnes, a 100-year storm that drastically reduced salinities and increased sedimentation throughout Chesapeake Bay (Hyer & Ruzecki 1976, Schubel 1976, Schubel et al. 1976), which resulted in a mass mortality event for *M. arenaria* (Cory & Redding 1976). Due to this storm, which followed declines in abundance of *M. arenaria* in the late 1960s, the commercial hydraulic dredge fishery for *M. arenaria* essentially ended in Virginia waters around 1968, and has failed to recover since then (Haven, 1970). More recent (post-1990) dramatic declines in abundance of *M. arenaria* have resulted in variable, low harvests in Maryland waters (Homer et al. 2011). Since 1980, commercial clammers have gradually switched to harvest of *T. plebeius* for eel and crab pot bait (Dungan et al. 2002, Homer et al. 2011). Like *M. arenaria, T. plebeius* populations have experienced declines in recent years, which were first documented in 2003 and resulted in the loss of 70-80% of the population in Maryland (Fig. 1; Homer et al. 2011). There are no historic landings records or long-term time series of *T. plebeius* abundance, so the history of decline and potential mechanisms for decline in this species are largely unknown.

**Figure 1.**
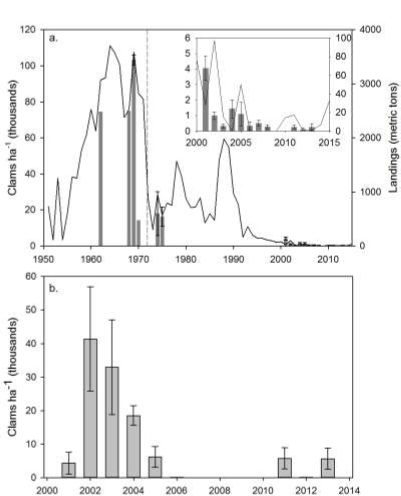
(a) *Mya arenaria* abundance (± S.E.) and fishery landings (solid line) for the period 1951-2015, with inset showing amplification of abundance and landings for 2000-2015. Abundance data (shaded bars) are from Maryland DNR (2001-2008) and SERC (2011-2013) fishery-independent escalator dredge sampling. Landings are for the Maryland and Virginia portions of Chesapeake Bay combined (NMFS Commercial Landings Database). Vertical dashed line represents Tropical Storm Agnes (1972). Data source: NMFS Commercial Landings Database. (b) *Tagelus plebeius* abundance (± S.E.) from Maryland DNR (2001-2008) and SERC (2011-2013) fishery-independent escalator dredge sampling. Note differences in y-axis ranges.

Multiple factors likely control densities of *M. arenaria* and *T. plebeius* in the Chesapeake Bay and other systems, including predation (Seitz et al. 2001, Beal 2006), overfishing (Brousseau 2005), low recruitment (Beukema & Dekker 2005, Bowen & Hunt 2009), rising temperatures (Najjar et al. 2000), habitat loss (Glaspie 2017), and disease mortalities (Dungan et al. 2002). Some of these factors, in particular disease and overharvesting, have been blamed for the inability of *M. arenaria* to recover from Tropical Storm Agnes and the recent declines in *M. arenaria* and *T. plebeius* abundances (Fisher et al. 2011, Homer et al. 2011). Overharvesting is unlikely to be responsible for current trends in Chesapeake Bay clam densities because, as previously mentioned, fishing pressure is not consistent throughout the Bay. The role of the remaining factors in determining local or regional population dynamics of *M. arenaria* and *T. plebeius* has yet to be determined.

There is some evidence that habitat preferences, including availability of refuge habitat that allows clams to avoid predation, may influence population dynamics of *M. arenaria* and *T. plebeius.* Habitat type has been shown to influence the distribution and abundances of *M. arenaria.* Growth rates can be impacted by sediment type, with higher growth rates observed in sand than mud-gravel-shell mixtures (Newell 1982). Habitats with more structure, such as gravel, shell hash, or seagrass, can also provide refuge from predators (Sponaugle & Lawton 1990, Skilleter 1994, Seitz et al. 2001). The availability of some refuge habitats, such as seagrass and oyster shell, is decreasing in the Chesapeake Bay (Rothschild et al. 1994, Lefcheck et al. 2017). The severe and persistent declines of Chesapeake Bay seagrass in the recent past have been attributed mostly to anthropogenic nutrient and sediment pollution, which decrease water clarity (Kemp et al. 2004). The dominant seagrass species in Chesapeake Bay, eelgrass *Zostera marina,* is also sensitive to warming, and the interaction between warming and poor water quality has resulted in recent declines of shallow beds (Lefcheck et al. 2017). In the past several years, seagrass die-offs induced by extreme high temperatures in Chesapeake Bay have resulted in the prediction that *Z. marina* may disappear from the Bay entirely (Moore & Jarvis 2008; though seagrass may be increasing in the coastal bays, see Orth & McGlathery 2012). Loss of oyster reefs in Chesapeake Bay due to overfishing, habitat destruction, and disease has resulted in the ecological extinction of oysters in the Bay (Rothschild et al. 1994, Wilberg et al. 2011). As oysters have been lost, so has a major source of shell material in the Bay. Oyster shell is a limited resource, and shell half-life is estimated to be as little as 3-10 years in Delaware Bay (Mann & Powell 2007). Loss of this structural refuge may have implications for populations of deep-burrowing clams such as *M. arenaria* and *T. plebeius.*

The recent findings of high prevalences of the parasitic protist *Perkinsus chesapeaki* in *M. arenaria* and *T. plebeius* suggest that this could be a cause of the population declines of both bivalve species (Dungan et al. 2002, Reece et al. 2008). *Perkinsus chesapeaki* infections have reached epizootic levels in *M. arenaria* (Farley et al. 1991, Dungan et al. 2002, Reece et al. 2008). However, prevalence is not necessarily equal to pathogenicity, as seen in some disease-resistant oyster stocks in the Chesapeake Bay (Encomio et al. 2005). The cancer disseminated neoplasia also causes mortality of *M. arenaria,* but this disease is not as prevalent as *P. chesapeaki,* and is not reported to affect *T. plebeius* (Dungan et al. 2002); thus, the disease aspect of this study focuses only on the impact of *P. chesapeaki* on *M. arenaria* and *T. plebeius.*

Extreme temperature, dissolved oxygen, or salinity levels may be stressful to *M. arenaria* and *T. plebeius,* resulting in mortality or metabolic constraints that make these conditions not suitable for growth. *Mya arenaria* is near the southern extent of its geographic range in Virginia, and is not tolerant of temperatures above 28°C, a temperature that is frequently exceeded in Chesapeake Bay during summer months (Moore & Jarvis 2008). In contrast, *T. plebeius* is a warm-water species that is distributed into South America (Abrahao et al. 2010), and high water temperatures (above 28°C) may not be stressful for this species. Benthic species in general are intolerant of dissolved oxygen levels below 1.5 mg L^−1^ (Rosenberg et al. 1991), and*M. arenaria* will decrease burial depth and extend their siphons into the water column under severe hypoxia (< 1.5 mg L^−1^) (Taylor & Eggleston 2000). Thus, low dissolved oxygen is likely to be stressful for both *M. arenaria* and *T. plebeius.* Finally, *T. plebeius* are rarely found at salinities under 5 ppt (Holland et al. 1987), and*M. arenaria* are intolerant of salinities below 4 ppt (Abraham & Dillon 1986).

Settlement and post-settlement processes are also important in determining distribution of organisms in soft-sediment communities (Olafsson et al. 1994). Larval behavior, hydrodynamics, and intense predation of postlarval bivalves control patterns of *M. arenaria* recruitment in other systems (Beukema & Dekker 2005, Bowen & Hunt 2009). Recruitment of *M. arenaria* remains high in several tributaries of Chesapeake Bay (Lovall et al. 2017) though these individuals rarely survive to adulthood, except in habitats with sufficient structure to allow protection from predators (Seitz et al. 2005). Locally, recruitment of *M. arenaria* is not necessarily correlated with abundance of adult clams (Bowen & Hunt 2009), but low densities of *M. arenaria* and *T. plebeius* throughout the Chesapeake Bay may limit recruitment and contribute to the loss of these bivalve species, as seen in bay scallops *Argopecten irradians irradians* in Long Island Sound (Tettelbach et al. 2015). In addition, larval and post-larval individuals of *M. arenaria* are more susceptible to extreme environmental conditions such as high temperatures and low salinities (Abraham & Dillon 1986), and population dynamics for this species may be controlled by this sensitive life stage.

No single factor can likely be exclusively attributed to the declines of these two bivalve species; thus, multiple factors were considered to gain useful insight into the problem. In this study, we used field surveys to concurrently examine predation, habitat, recruitment, and disease prevalence/intensity to disentangle the relative effects of these factors on survival and persistence of *M. arenaria* and *T. plebeius.* We hypothesized that: (1) densities of *M. arenaria* and *T. plebeius* are positively associated with presence of complex habitat such as seagrass and shell, and negatively correlated with predator densities; (2) densities of *M. arenaria* are negatively correlated with temperature; and (3) intensity and prevalence of infection by presumed *Perkinsus chesapeaki* (hereafter, *Perkinsus* sp.) increase under stressful environmental conditions (i.e., positively correlated with temperature and river discharge, negatively correlated with salinity and dissolved oxygen).

## MATERIALS AND METHODS

### Study system

The Chesapeake Bay provides an ideal location to study the effects of predators, habitat type, recruitment, disease, and physical factors on the distribution and abundance of *M. arenaria* and *T. plebeius.* The Bay offers a range of environmental conditions including temperature, salinity, dissolved oxygen, and availability of complex habitat. Lower Chesapeake Bay (the Virginia portion of Chesapeake Bay) is mostly polyhaline (except in the upper reaches of the tributaries). Upper Chesapeake Bay (the Maryland portion of the Bay) is mostly mesohaline. Bottom habitat type in the upper Bay is characterized by oyster shell hash, soft muds, fine sands, and gravel/pebbles (Smith et al. 2003). Bottom type in the lower Bay is similar to that of the upper Bay (Wright et al. 1987), except shallow shoals in the lower Bay south of the Potomac River often have mixed beds of eelgrass *Z. marina* and widgeongrass *Ruppia maritima* (Orth et al. 2010). The entire Chesapeake Bay experiences seasonal hypoxia in deep channel water, which occasionally wells up onto the shoals with the tides and wind events (Sanford 1990, Kemp et al. 2005). Summer temperatures in the upper Bay are likely still hospitable for *M. arenaria,* while the lower Bay experiences periods when they are above the tolerance limit for the species.

In Chesapeake Bay, the dominant predators of *M. arenaria* and *T. plebeius* include the blue crab *Callinectes sapidus,* horseshoe crabs *Limulus polyphemus* (Botton 1984, Lee 2010), and demersal fishes (de Goeij et al. 2001, Seitz et al. 2001) including cownose rays *Rhinoptera bonasus* (Fisher 2010). Crabs forage for clams from the sediment and consume the entire clam (Beal 2006), whereas demersal fishes nip clam siphons, causing clams to reduce their burial depth and exposing them to increased predation by probing predators (de Goeij et al. 2001). High predation rates on infauna are also associated with seasonal migratory behavior of cownose rays (Blaylock 1993), which are able to consume bivalves that would otherwise avoid predation by burrowing, armor, and/or size refuges (Fisher 2010).

### Survey design

Clams were collected from three subestuaries of lower Chesapeake Bay (Lynnhaven River, York River, Mobjack Bay), and three subestuaries of upper Chesapeake Bay (Western Shore, Eastern Bay, Chester River) in fall 2011; spring/summer/fall 2012; and either spring/summer 2013 (for lower Bay) or summer/fall 2013 (for upper Bay; see Table 1 for sampling dates). Each sampling season, four to nine sites within each subestuary were sampled; sites were specific tributaries or shorelines in the subestuary that were chosen to represent the range of available benthic habitat types (mud, sand, gravel, shell, or seagrass) available in the subestuary (Table 1).

**Table 1.**
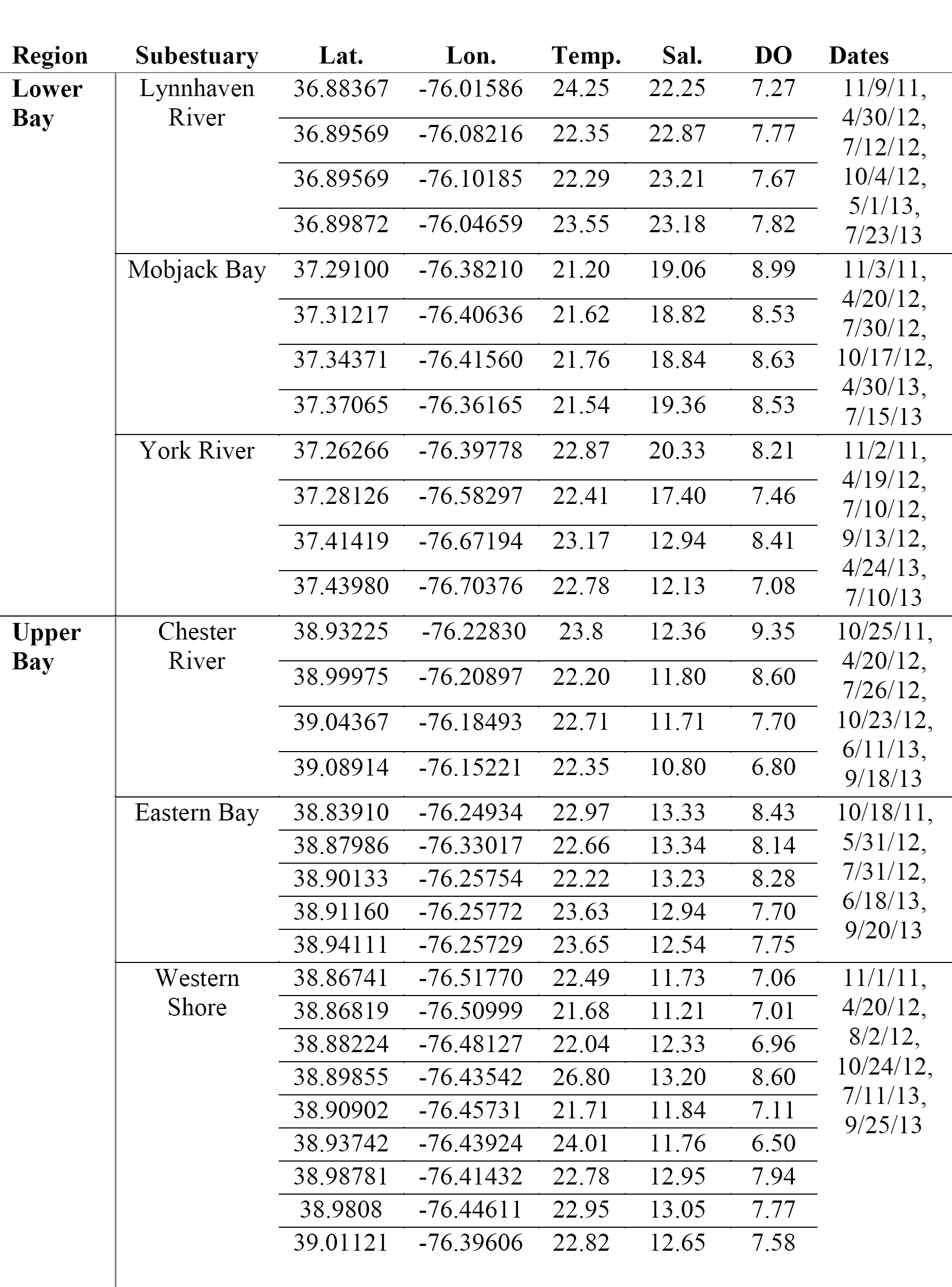
Locations and dates of suction sampling in the lower Chesapeake Bay and hydraulic dredge sampling in the Upper Chesapeake Bay. Sites (latitude = Lat. and longitude = Lon.) were each sampled on multiple dates, listed here for each subestuary. Means are shown for temperature (Temp., °C), salinity (Sal.) and dissolved oxygen (DO, mg L^−1^).

At each sampling site in the lower Bay, three replicate samples were collected in each season from shallow water of 1.5-2 m depth mean high water using a suction sampling device that collects samples of 0.11 m^2^ area and 40 cm depth. Replicate sample locations in each site were selected by throwing the suction cylinder from the boat, and samples were sieved through 3-mm mesh. All samples were assigned a habitat category (mud, sand, gravel, shell, or seagrass) based on observations made in the field and the lab during sample processing. In the upper Bay, due to the low density of clams and a lack of seagrass in clam habitat, samples were collected using a commercial hydraulic escalator dredge targeting areas fished for *M. arenaria.* This method of collection was not used in the lower Bay to prevent unnecessary destruction of existing seagrass beds, and because clams were consistently captured using less destructive methods such as suction sampling. Habitats were categorized by the presence/absence of oyster shell hash retained on the escalator dredge, and by diver surveys conducted in 2012. All *M. arenaria* and *T. plebeius* were counted, and clam densities at each site were determined by sample area (in lower Bay) or dredge distance (in upper Bay). All clams were measured to the nearest 1 mm for shell length. Bottom (e.g. < 1 m from the sediment surface) dissolved oxygen, salinity, and temperature were also recorded at each site using a YSI (Model 85, Yellow Springs Instruments).

Between spring 2012 and summer 2013, blue crab abundance was quantified at each lower Bay site within a few days of bivalve suction sampling using six replicate 20-m tows of a modified commercial crab scrape (usually used for harvesting soft-shell peeler crabs in seagrass in lower Chesapeake Bay; 6 mm mesh, 1 m width; Seitz et al., 2008) because it is effective sampling mobile fauna in seagrass. In the upper Bay, seagrass was not a concern and two replicate 4.9 m-wide otter trawl tows were conducted at each site in the spring, summer, and fall of 2012 for seven minutes each (approx.∼ 500 m). Gear efficiencies of both methods are similar: 24-34% for capture of juvenile crabs with the crab scrape (Ralph 2014) and ∼22% for capture of most predators, including fish and crabs, with the otter trawl (Homer et al. 1980). Any fish caught in tows were counted and released. A fish species list and number caught can be found in Supplemental Table 1. Approximately 10% of bivalve sampling events were paired with trawls that occurred approximately a month before or after bivalve sampling, due to logistical constraints. In this case, an average of crab or fish abundance before and after the bivalve sampling was used in analyses. At each bivalve sampling site and for each season, the number of ray pits within 1 m to either side of a 50-m transect were counted, and are treated as a proxy of cownose ray density (Hines et al. 1997). Ray pits were about 0.3 m in diameter and 10 cm deep, and could be easily seen in good visibility, or detected by sweeping the sediment with hands in poor visibility. Horseshoe crabs were not quantified because they were not abundant enough in any of our samples. Fish and crab abundances per tow and ray pit counts were converted to density per 100 m^2^ for analyses. Due to logistical constraints, certain data, such as environmental data and predator abundance, were missing from the survey dataset, especially for fall 2011, so models that included these missing variables were fit to data from spring 2012 through fall 2013. Environmental data from the CBIBS monitoring buoy at 38.963N 76.448W were used for Maryland western shore samples from summer and fall 2013.

We conducted a gear comparison of bivalve sampling methods by using suction samples collected in the same areas as hydraulic dredging in spring 2012. Site-average densities from suction samples were regressed against site-average densities from hydraulic dredging for both *M. arenaria* and *T. plebeius. Mya arenaria* densities calculated using suction sampling were 3.58 times higher than those calculated from hydraulic dredge samples (R^2^ = 0.95; Supplementary Fig. 1a); thus, suction sample densities for *M. arenaria* were reduced by a factor of 3.58 to allow both data sets to be analyzed together. Densities calculated for *T. plebeius* using both methods were in close agreement (slope = 1.10, R^2^ = 0.90; Supplementary Fig. 1b), so the raw data sets were combined for analyses. Suction samples appeared to collect more small and large *Mya arenaria* (mean 42.33 mm, SD 13.73 mm; Supplementary Fig. 2a) than dredge sampling (mean 40.22 mm, SD 5.42 mm; Supplementary Fig. 2b), indicating that the discrepancy in densities was not an artifact of hydraulic dredge mesh size. Biomass of both *M. arenaria* and *T. plebeius* from the lower Bay was determined as ash free dry weight of bivalves dried in a drying oven for 24 hours, and ashed in a muffle furnace at 550 °C for five hours; bivalves in the upper Bay were not processed for biomass.

The long-term trend of *M. arenaria* recruitment in Maryland was evaluated using data from surveys conducted by SERC in the Rhode River, Maryland. Benthic core samples were collected approximately quarterly (typically March/April, June, October, December) at two sandy subtidal sites from 1981-2016. These sites were located at 38.885797N, 76.541790W and 38.868377N, 76.518276W. Cores were 10.2 cm in diameter and 35 cm long. Seven core samples were taken per site on each sample date and animals retained on a 500 μm sieve were preserved in 10% buffered formalin and stained with rose Bengal. All *M. arenaria* were measured for shell length and individuals <10 mm shell length were considered recruits. The total number of recruits in each year was calculated by site and then averaged across the two sites to provide an index of annual recruitment.

### Disease

Whenever possible, 30 clams from each upper Bay site were held for 24-96 h in flow-through systems of ambient Rhode River or Tred Avon River waters before they were examined for *Perkinsus* sp. infections. Clams were dissected to secure labial palp tissues that were inoculated into tubes containing 3 mL of Ray’s fluid thioglycollate medium (RFTM, Ray 1966), and then incubated in the dark at 28 °C for 4-7 d. After incubation in RFTM, individual clam tissues were macerated in pools of 25% Lugol’s iodine before microscopic examinations at 40x magnification. *Perkinsus* sp. hypnospores were quantified in the most heavily infected palps of individual clams, and infections were assigned ordinal intensity ranks of 0-5.

*Perkinsus* sp. prevalence and intensity data for 2011-2013 were analyzed together with similar data collected 2000-2009. Only data from clams collected at the same geographic locations by both investigations were used for the time series. For samples collected during our 2011-2013 survey, infection intensity categories included the rank of 0.5 for infections of very low intensities (Ray 1954). In accordance with suggested method for cross-calibrating such data (Dungan & Bushek 2015), infection intensity data for 2011-2013 were cross calibrated to those of 2000-2009 by pooling infection intensities of the lowest ranks (0.5, 1.0) (Dungan et al. 2002, Reece et al. 2008, Homer et al. 2011). Prevalences were estimated as the proportion of infected clams.

Clams tested for *Perkinsus* sp. infections were predominantly collected from upper Bay sites where hydraulic dredging was used to reliably collect adequate numbers of adult clams. Although hydraulic escalator dredges were not used in the lower Bay, 30 juvenile *M. arenaria* were collected by hand from Indian Field Creek, York River, VA, in April 2013, held in flow-through tanks for several months until they were 20-30 mm in shell length, and were assayed for *Perkinsus* sp. infections in July 2013.

### Statistical analyses

Both *M. arenaria* and *T. plebeius* exhibited many instances of zero catch, so two models were used to analyze the data: presence/absence was modeled with a binomial generalized additive model (GAM, logit link), and non-zero densities were modeled with a Gaussian GAM (identity link) on log-transformed data. GAMs for *T. plebeius* included the following as parametric predictors: temperature (°C), salinity, dissolved oxygen (mg L^−1^), ray pit density (m^−2^), crab density (m^−2^), fish density (m^−2^), and habitat (5 categories: gravel, mud, sand, shell, and seagrass). *Mya arenaria* were rarer and data contained many more zeroes than *T. plebeius*; to allow for model convergence, avoid over-smoothing, and ensure homoscedasticity of residuals, habitat was reduced to two categories (simple, including mud, sand, and gravel; and complex, including seagrass and shell) in both presence/absence and non-zero density models, and the presence/absence model was run only on spring/summer data for 2012. *Perkinsus* sp. infection intensity rank-score was modeled as a log-transformed continuous variable using a Gaussian GAM (identity link). Disease GAMs for *M. arenaria* and *T. plebeius* included the following predictors: temperature (°C), salinity, dissolved oxygen (mg L^−1^), bivalve length (mm), and bivalve density (m^−2^). To account for trends in data across space, a two-dimensional spline (trend-surface) was fit on latitude and longitude for all GAMs (Cressie 1993). To account for trends in time, sampling day was fit in each model using a spline with a kernel smoothing function (Kohn et al. 2002). The number of knots (k) in each model was chosen automatically using the Generalized Cross-validation (GCV) optimizer in the R package ‘mgcv’ (Wood 2017), except for the presence/absence model for *M. arenaria,* which was fit with a spline of order k = 2 because only two seasons were modeled and the GCV optimizer produced a model with skewed residuals.

All variables were examined for multicollinearity with scatter plots and Pearson correlation coefficients before inclusion in the model. There was no evidence of multicollinearity; all variable combinations had Pearson correlation coefficients ≤ 0.8 (Berry & Feldman 1985; Supplementary Figure 3). For all GAMs, appropriateness of the smoothing parameter was assessed using the k-index method and the smoothing parameter chosen was deemed adequate if the simulated p-value was > 0.05 (Wood 2017). Homoscedasticity of residuals was assessed visually using residual and quantile-quantile plots. All models had appropriate smoothing and met the assumption of homogeneity of variance.

Time series (infection intensity, infection prevalence, and *M. arenaria* recruitment) were analyzed using automatic time series forecasting (ARIMA models; Hyndman & Khandakar 2008), and disease models included spring/summer temperature (April-August) obtained from the Chesapeake Bay Program Water Quality Database (http://www.chesapeakebay.net/data) for all tidal mainstem Chesapeake Bay stations, and annually averaged Susquehanna River discharge obtained from the USGS water quality monitoring station at Harrisburg, PA (National Water Information System at https://waterdata.usgs.gov/usa/nwis). No autocorrelation or non-stationarity was detected in ARIMA models of *Perkinsus* sp. infection intensities or prevalences ([p,d,q] = [0,0,0]), so analysis proceeded using linear models. For multiple comparisons, significant difference was determined using non-parametric bootstrap hypothesis testing with 10,000 simulations and α = 0.05 (Efron & Tibshirani 1993), and Cohen’s d was calculated as a measure of effect size for all two-group comparisons. All analyses were completed in R (R Core Team 2017), and code and data files have been provided as supplementary material.

## RESULTS

### Survey

Temperature over the course of the survey (fall 2011 – summer 2013) ranged 12.6 °C – 33.3 °C. Temperatures exceeding 28 °C and 30 °C were observed for 23.7% and 6.5% of all samples collected, respectively. Salinity ranged from 5.3 psu in the upper York River site to 25.7 psu in the Lynnhaven Bay sites. The minimum dissolved oxygen level observed was 3.3 mg L^−1^ in Mobjack Bay, though the site means ranged 6.50 mg L^−1^ – 9.35 mg L^−1^ (Table 1).

Densities of *M. arenaria* were 2.75 times higher at sites in the lower Bay than at sites in the upper Bay (p = 0.02, d = 0.01, Fig. 2a). The maximum density observed was 53.33 m^−2^ *M. arenaria* near the mouth of the York River, VA, in spring 2012. Out of all samples collected, 54.73% of upper Bay and 89.82% of lower Bay samples did not contain *M. arenaria.* Despite repeated sampling, samples from four sites in Virginia contained no *M. arenaria* (two in Mobjack Bay and two in Lynnhaven). Throughout all samples, densities of *Tagelus plebeius* were 13.07 times higher than *M. arenaria* densities (p < 0.001, d = 0.02, Fig. 2b). Densities of *Tagelus plebeius* were 8.37 times higher at sites in the lower Bay than at sites in the upper Bay (p < 0.001, d = 0.04). The maximum density observed for *T. plebeius* was 218.18 m^−2^ in the lower Bay in Lynnhaven (Linkhorn Bay) in summer 2013. *Tagelus plebeius* were found at every lower Bay site, but were not found at two sites in the Eastern Bay of Maryland throughout the study period. Trends in biomass of both *M. arenaria* and *T. plebeius* in the lower Bay closely matched trends in densities (Supplementary Figure 4).

**Figure 2.**
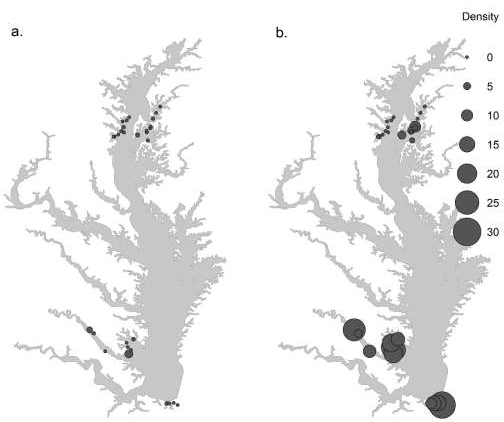
Mean density (m^−2^) of (a) *Mya arenaria* and (b) *Tagelus plebeius* captured in suction and hydraulic dredge samples between fall 2011 and summer 2013. Point size is a linear function of mean density at each site: Radius = (0.25 × Density) + 1

Average shell length of *M. arenaria* was 1.43 times greater at sites in the upper Bay than at sites in the lower Bay (p < 0.001, d = 0.07; Fig. 3a). Similarly, *T. plebeius* at sites in the upper Bay were 2.07 times larger than those at sites in the lower Bay (p < 0.001, d = 0.04; Fig. 3b). Only 5.56% of *M. arenaria* collected in the lower Bay were greater than 50 mm in shell length, while 29.73% of upper Bay *M. arenaria* were 50 mm or larger.

**Figure 3.**
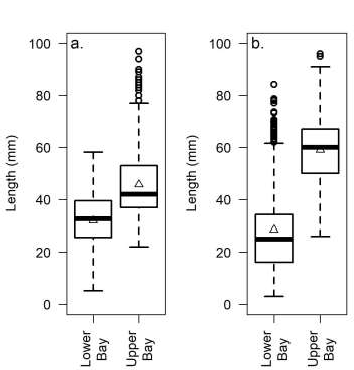
Average shell length in lower and upper Chesapeake Bay for (a) *Mya arenaria* and (b) *Tagelus plebeius,* with boxes from the first to third quartile, a horizontal line at the median, and a triangle at the mean. Whiskers extend from the lowest data point that is still within 1.5 inter-quartile range (IQR) of the lower quartile, to the highest data point still within 1.5 IQR of the upper quartile, and data outside this range (outliers) are shown as circle symbols.

The highest densities of *Mya arenaria* each year occurred in the spring, with declining density through the fall (Fig. 4a). *Tagelus plebeius* densities did not show consistent seasonal trends over the study period. In the lower Bay, *T. plebeius* densities were the lowest in spring 2012, increased through the summer and fall, and then remained high (generally 15-20 m^−2^) and stable through 2013 (Fig. 4b). In contrast, in the upper Bay the clearest seasonal trend involved a pulse of *T. plebeius* in spring 2012 (∼5 m^−2^) that disappeared through the summer and fall (Fig. 4a).

**Figure 4.**
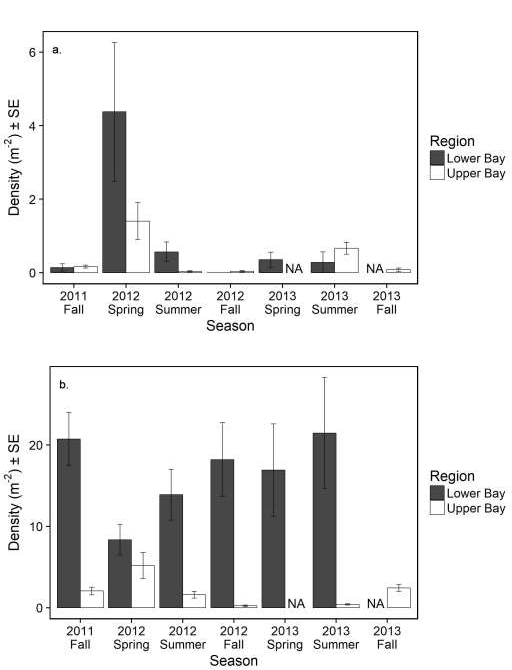
Mean densities and standard errors for (a) *Mya arenaria* and (b) *Tagelus plebeius* from fall 2011 through fall 2013 for the lower Bay (black) and upper Bay (white). NA = not available (not sampled). For the lower Bay, N = 36 for all seasons. For the upper Bay, N = 62, 51, 67, 59, NA, 43, 67 for Fall 2001 to Fall 2013, respectively.

Density of *M. arenaria* was higher in complex habitat (seagrass and shell) than in less complex habitat (mud, sand, and gravel) (Fig. 5a). The odds of finding *M. arenaria* in seagrass or shell were 392.04 times greater than in less-complex habitats such as mud, sand, and gravel. When *M. arenaria* were present, densities in seagrass or shell were 2.15 times greater than in less-complex habitats such as mud, sand, and gravel. The odds of finding *M. arenaria* decreased as temperature increased, and when they were present, *M. arenaria* density was negatively correlated with dissolved oxygen (Table 2). There was no significant relationship between *M. arenaria* density and predator density (Table 2).

**Figure 5.**
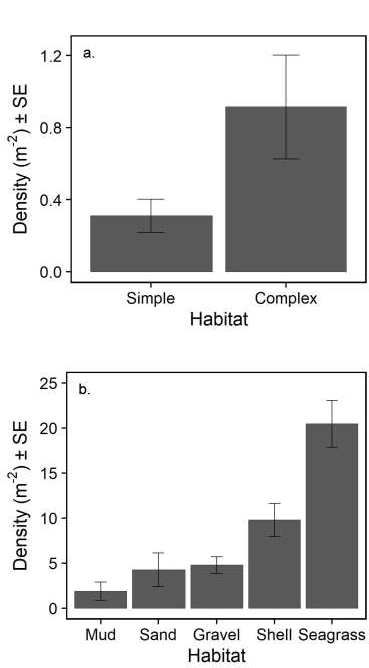
Trends in density for (a) *Mya arenaria* and (b) *Tagelus plebeius* in different habitats. Error bars denote standard error. For *M. arenaria,* N = 314 and 251 for simple and complex habitat, respectively. For *T. plebeius,* N = 57, 32, 182, 187, 64 for mud, sand, gravel, shell, and seagrass, respectively.

**Table 2.** Generalized additive model (GAM) coefficients and standard deviations (in parentheses) for presence/absence (P/A), density, *Perkinsus* sp. infection intensity (average ordinal intensity rank of infected individuals, scale 0-5) and prevalence (Dis. Prevalence) of infection (proportion of individuals infected) models for *Mya arenaria* and *Tagelus plebeius.* Transformations (either log or logit) are indicated next to the variable name and coefficients are not back transformed. Parametric predictor variables were temperature (temp., °C), salinity (Sal.), dissolved oxygen (DO, mg L^−1^), ray pit density (rays m^−2^), crab density (crabs m^−2^), fish density (fish m^−2^), and habitat (gravel [GR], mud [MU], sand [SA], shell [SH], and seagrass [included in the intercept, int.]). Disease GAMs also included bivalve length (mm), and bivalve density (m^−2^). Coefficients significant at α = 0.05 are bolded. N = number of observations used to fit each model.

|  | *Mya arenaria* |  |  |  | *Tagelus plebeius* |  |  |  |
| --- | --- | --- | --- | --- | --- | --- | --- | --- |
|  | logit(P/A) | log(Density) | log(Intensity) | Dis. Prevalence | logit(P/A) | log(Density) | log(Intensity) | Dis. Prevalence |
| *Int.* | **147.01 (62.41)** | 6.11 (7.09) | **-21.90 (2.09)** | **-23.01 (2.10)** | **12.14 (5.45)** | -4.56 (4.22) | **-4.23 (1.73)** | -3.18 (1.70) |
| *Temp.* | **-4.79 (2.34)** | -0.27 (0.29) | **0.44 (0.12)** | **0.60 (0.07)** | -0.28 (0.16) | 0.01 (0.11) | **0.19 (0.07)** | **0.14 (0.07)** |
| *Sal.* | -1.27 (0.75) | 0.17 (0.17) | **0.27 (0.13)** | **0.48 (0.04)** | -0.29 (0.16) | **0.31 (0.14)** | **-0.18 (0.06)** | 0.06 (0.05) |
| *DO* | -0.55 (0.78) | **-0.32 (0.16)** | **0.54 (0.06)** | **0.19 (0.03)** | 0.22 (0.17) | -0.04 (0.1) | -0.03 (0.03) | **0.10 (0.03)** |
| *Rays* | -7.94 (15.51) | 0.09 (1.76) | NA | NA | 1.45 (2.89) | 0.63 (1.25) | NA | NA |
| *Crabs* | 0.34 (0.77) | -0.04 (0.37) | NA | NA | -0.09 (0.18) | -0.01 (0.15) | NA | NA |
| *Fish* | -0.37 (0.65) | 0.10 (0.17) | NA | NA | -0.12 (0.15) | -0.15 (0.12) | NA | NA |
| *Habitat* | SG & SH v. | SG & SH v. | NA | NA | GR: -0.46 (1.32) | GR: 0.15 (0.97) | NA | NA |
|  | MU, SA & GR | MU, SA & GR |  |  | MU: -2.46 (1.48) | MU: 1.27 (0.96) | NA | NA |
|  | **-5.97 (2.82)** | **-0.77 (0.38)** |  |  | SA: -1.37 (0.81) | SA: **1.68 (0.65)** | NA | NA |
|  |  |  |  |  | SH: **-2.42 (0.69)** | SH: 0.13 (0.59) | NA | NA |
| *Bivalve length* | NA | NA | **0.06 (0.01)** | **2.89x10<sup>-2</sup> (4.01x10<sup>-3</sup>)** | NA | NA | **0.04 (0.01)** | -0.02 (0.01) |
| *Bivalve density* | NA | NA | 8.13x10<sup>-5</sup> (2.26x10<sup>-3</sup>) | 4.11x10<sup>-4</sup> (1.62x10<sup>-3</sup>) | NA | NA | 1.78x10<sup>-3</sup> (3.52x10<sup>-3</sup>) | 1.04x10<sup>-3</sup> (2.92x10<sup>-3</sup>) |
| *N* | 150 | 62 | 102 | 88 | 310 | 179 | 154 | 128 |

*Tagelus plebeius* density in seagrass was higher than in less-complex habitats (shell, gravel, sand, and mud) (Fig. 5b). The odds of finding *T. plebeius* in seagrass were 11.65 times greater than in mud (z = 1.48, p = 0.10), 3.95 times greater than in sand (z = 0.81, p = 0.09), and 11.28 times greater than in shell (z = 0.69, p = 0.0005). When *T. plebeius* were present, densities in sand were 4.63 times greater than in gravel, 5.39 times greater than in seagrass, and 4.74 times greater than in shell. High densities in sand were only observed in the spring (Fig. 5b). High densities of *T. plebeius* were also observed in shell, though this was not significant in the models (Table 2). When they were present, *T. plebeius* density increased with salinity (Table 2). There was no significant relationship between *T. plebeius* density and temperature, dissolved oxygen, or predator density (Table 2).

*Mya arenaria* recruitment in the Rhode River declined steadily between 1980 and 2016. Recruitment was characterized by an Autoregressive Integrated Moving Average (ARIMA[2, 1, 1]), with a negative trend (drift) of -0.03 million clams ha^−1^ y^−1^. The survey captured no recruits post-2005 (Fig. 6).

**Figure 6.**
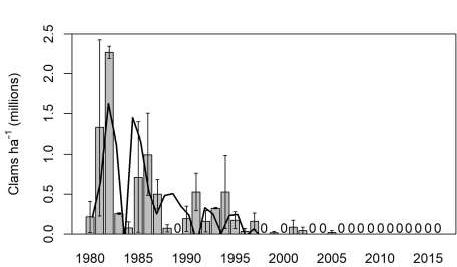
Mean annual *Mya arenaria* recruitment from fishery-independent core sampling in the Rhode River, Maryland (1980-2016). Recruitment data include only clams ≤ 10 mm shell length. Shown are means of two sampling sites (bars), standard error (error bars) and fit from ARIMA model (solid black line). Years where sampling occurred but no *M. arenaria* were captured are denoted by a “0”; no clams were captured in samples after 2005.

### Disease

Mean infection intensity for *Mya arenaria* was light at upper Bay sites, with a mean of 0.48 on a 5-point scale (Fig. 7a). Infection intensity in *M. arenaria* increased by a factor of 0.55 for each degree increase in temperature, and increased by a factor of 0.07 for each mm in shell length (back-transformed from Table 2). *Mya arenaria* infection intensity was positively correlated with temperature, salinity, and dissolved oxygen (Table 2). The mean infection intensity for *T. plebeius* was light to moderate and was 1.57 times greater than that for *M. arenaria* (p < 0.001, d = 0.03; Fig. 7b). Mean infection intensity for *T. plebeius* decreased by a factor of 0.16 with each unit increase in salinity, and increased by a factor of 0.21 with each unit increase in temperature (back-transformed from Table 2). Infection intensity increased with *T. plebeius* size, but did not increase with bivalve density for either *M. arenaria* or *T. plebeius* (Table 2).

**Figure 7.**
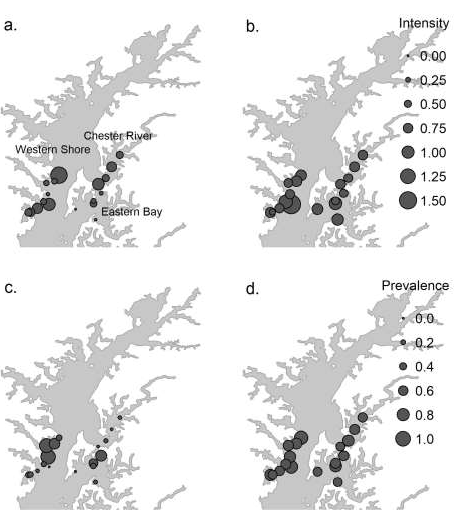
Average *Perkinsus* sp. (a) infection intensity (average ordinal intensity rank of individuals 0-5) for *Mya arenaria,* (b) infection intensity for *Tagelus plebeius,* (c) prevalence of infection (proportion of individuals infected) for *M. arenaria,* and (d) prevalence of infection for *T. plebeius* for the upper Chesapeake Bay from hydraulic dredge samples collected between fall 2011 and fall 2013. Point size is a linear function of mean disease intensity (Radius = [2.0 × Intensity] + 0.5) or mean disease prevalence (Radius = [2.5 × Prevalence] + 0.5) at each site.

*Perkinsus* sp. infection prevalence was low for *M. arenaria,* averaging 29.21% over the course of the survey (2011-2013). *Perkinsus* sp. infection prevalence during the survey (2011-2013) was 2.17 times higher in *T. plebeius* (Fig. 7c) than in *M. arenaria* (Fig. 7d; p < 0.001, d = 0.06). Mean infection prevalence for both bivalve species increased with temperature and dissolved oxygen, and prevalence also increased with salinity for *M. arenaria* (Table 2).

Infection prevalence increased with bivalve size for *M. arenaria,* but not for *T. plebeius* (Table 2). Infection prevalence and intensity were not related to *M. arenaria* or *T. plebeius* densities (Table 2). In the lower Bay, where 30 juvenile *M. arenaria* were assayed, hypnospores were present in two individuals at very low infection intensities (ordinal rank 0.05).

Prevalence of infection was moderate to high between 2000 and 2013, with mean infection rates of 68.46% for *M. arenaria* and 81.86% for *T. plebeius* (Fig. 8a). Across all years in the time series, average prevalence of infection was similar between species (d = 0.25, p = 0.16), and average infection intensities were light and similar between *M. arenaria* and *T. plebeius* (d = 0.04, p = 0.18). Infection prevalence and intensity were negatively correlated with annual Susquehanna River discharge for*M. arenaria* (prevalence: coef. = −1.25×10^−5^, t_9_ = −3.40, p = 0.01; intensity: coef. = −2.28 ×10 ^−5^, t_9_ = −2.31, p = 0.05) but not for *T. plebeius* (prevalence: coef. = −6.99×10^−6^, t_8_ = −1.60, p = 0.15; intensity: coef. = −1.17×10^−5^, t_8_ = −1.28, p = 0.24; Fig. 8c). Infection prevalence and intensity were not correlated with spring/summer temperature for *M. arenaria* (prevalence: coef. = 2.96×10^−2^, t_9_ = −0.56, p = 0.59; intensity: coef. = 6.23×10^−2^, t_9_ = 0.44, p = 0.67) or for *T. plebeius* (prevalence: coef. = −2.51×10^−2^, t_8_ = −0.40, p = 0.70; intensity: coef. = −9.95×10^−2^, t_8_ = −0.76, p = 0.47).

**Figure 8.**
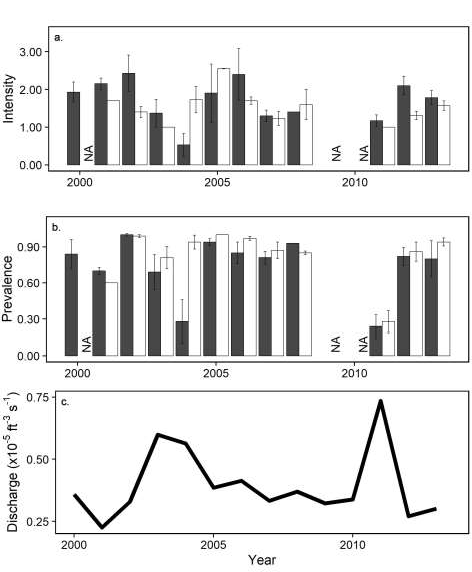
Time series of *Perkinsus* sp. (a) mean annual infection intensity (average ordinal intensity rank) and (b) mean annual prevalence (proportion of individuals infected) for *Mya arenaria* (black) and *Tagelus plebeius* (white) from 2000-2013. Error bars indicate standard error. Also shown is (c) average annual Susquehanna River discharge as a proxy for salinity (from the USGS National Water Information System). NA = not available (not sampled). See supplementary data files for sample sizes.

## DISCUSSION

As we hypothesized, the distributions of thin-shelled commercial species *Mya arenaria* and *Tagelus plebeius* were positively associated with complex habitat. Densities of both *M. arenaria* and *T. plebeius* were associated with habitats with a high degree of complexity (seagrass and shell) as compared to some less-complex habitats (mud, sand, and gravel). Complex habitats may be more favorable for these species because they increase rates of larval settlement by baffling water currents (Heiss et al. 2010), provide increased food resources for both suspension and facultative deposit-feeding species (Peterson et al. 1984), and provide refuge from predators (Orth et al. 1984). However, this also implies that habitat loss may be an important factor in the decline of *M. arenaria* and *T. plebeius,* as both seagrass and oyster reef habitats are declining in Chesapeake Bay (Orth et al. 1984, 2006, Rothschild et al. 1994, Beck et al. 2011, Lefcheck et al. 2017).

In agreement with our hypotheses, densities of *M. arenaria* were negatively associated with increasing temperature. *Mya arenaria* is a cold-water species that is distributed from the sub-arctic regions to North Carolina along Atlantic coasts of North America (Abraham & Dillon 1986, Maximovich & Guerassimova 2003). Typically *M. arenaria* survives well in temperatures from 2 to 28 °C (Cohen 2005), with mortality usually occurring above 30 °C (Kennedy & Mihursky 1971). It is expected that with global climate change, Chesapeake Bay may become inhospitable for this species. The upper tolerance range for *M. arenaria* is frequently surpassed in the summer in Chesapeake Bay, especially in shallow waters where this sampling effort took place. High summer temperatures in particular are likely a factor in the decline of *M. arenaria* in lower Chesapeake Bay.

Bivalve presence/absence and density were not negatively associated with ray, blue crab, or fish densities for either species, in contrast to our hypothesis, and horseshoe crabs were not abundant enough to determine their relative contribution to bivalve mortality. However, the snapshot in time provided by our predator and bivalve sampling may not reflect the legacy of predator-prey interactions experienced by local clam populations (Thrush et al. 1997, Seitz & Lipcius 2001, Seitz et al. 2016). Our seasonal time series for *M. arenaria* abundance in particular seem to indicate that predation is an important factor in determining density. In both the upper and lower Bay, densities of *M. arenaria* decline throughout the summer months when predation peaks (Hines et al. 1990). Observed temporal trends also correspond with *M. arenaria* reproductive behavior in Chesapeake Bay, where the fall spawn is more successful than the spring spawn, due to decreased predation pressure in the winter months (Blundon & Kennedy 1982a, Baker & Mann 1991). Individuals spawned in the fall are able to settle and grow throughout the winter, when risk of predation is minimal, and this new generation manifests as a springtime spike in density. However, even the clams that survive until the spring are almost completely consumed by predators each year, essentially resulting in an “annual crop” rather than a stable population with a sustainable age distribution. The declines in *M. arenaria* in summer may be associated with increasing temperature as well as predation, and we have insufficient evidence from our snapshot in time to distinguish between effects of the two. However, this effect persisted in both the upper and lower Bay, and under different temperature regimes, indicating that predation is likely a major driver of observed trends in *M. arenaria* density.

Despite occupying a similar niche in Chesapeake Bay, *M. arenaria* and *T. plebeius* exhibited different seasonal trends in densities. *Tagelus plebeius* spawns in the spring (Holland & Dean 1977, da Silva et al. 2015), and while a spring high-density event was noted in the upper Bay in 2012, the lower Bay had relatively steady densities that did not show a clear recruitment signal or summer crash. *Tagelus plebeius* in the lower Bay likely do not exhibit the same seasonal crashes in abundance observed for *M. arenaria* because a robust adult *T. plebeius* population remains in lower Chesapeake Bay throughout the seasons. This population allows for high densities of many different size classes to exist at any given time. However, in the upper Bay, where densities are much lower, seasonal spring recruitment was evident in time series, indicating very different population dynamics in these two regions of Chesapeake Bay. *Tagelus plebeius* in the upper Bay seem to exhibit annual population fluctuations more similar to those of *M. arenaria.* This may be because of density-dependent mortality sources specific to the upper Bay, including fishing pressure and disease, which work to keep upper Bay *T. plebeius* at low densities that are sensitive to annual recruitment and predation.

Predation and low fecundity likely drive patterns in recruitment of *M. arenaria* in Chesapeake Bay, which is characterized by very low recruitment with some pockets of locally high recruitment, though studies examining recruitment in more locations would be necessary to identify factors contributing to low recruitment. Although there are some cases of locally high recruitment of *M. arenaria* in Virginia (Lovall et al. 2017), recruitment in the Rhode River, Maryland, declined drastically since the 1980s and now is essentially zero. Predation has been identified as the most important limiting factor in recruitment of *M. arenaria* in other regions (Beal et al. 2001, Hunt & Mullineaux 2002, Beal 2006), and likely plays an important role in Chesapeake Bay as well. In addition, low adult densities and small body sizes may limit recruitment, as fecundity increases exponentially with clam size (Brousseau 1978, Brousseau & Baglivo 1988).

In agreement with our hypothesis, *Perkinsus* sp. infection intensity and prevalence in both species was positively associated with high temperatures, conditions that were considered stressful for *M. arenaria* in particular. Impacts of dissolved oxygen on *Perkinsus* sp. infection were less clear, though dissolved oxygen was generally within the tolerable range for *M. arenaria* and *T. plebeius* (> 1.5 mg L^-1^; Rosenberg et al. 1991, Taylor & Eggleston 2000), and may not have been extreme enough to induce a stress response during the course of the survey. Disease intensity and prevalence in *M. arenaria* was positively correlated with salinity, and long-term intensities and prevalences of *Perkinsus* sp. infections in *M. arenaria* were negatively associated with Susquehanna River discharge, a proxy of salinity. Salinity preference of *Perkinsus* sp. may explain the long-term correlation between infection intensities/prevalences and river discharge/salinity. *Perkinsus* sp. survives and proliferates best at moderate to high salinities, depending on the species. At 28 °C, *Perkinsus chesapeaki* survival and proliferation is optimal at moderate salinities 15-25%_0_ (La Peyre et al. 2006) or 14 *%%* (McLaughlin et al. 2000). In oysters, infection intensity and prevalence of a similar protist, *Perkinsus marinus,* were both greater in higher salinity waters (Burreson & Calvo 1996). Thus, it may be expected that *Perkinsus* sp. infection in *M. arenaria* would be greatest in high-salinity (low flow) years and locations.

It is possible that disease dynamics do not drive trends in *M. arenaria* and *T. plebeius* density in space and time, as intensities of infections were generally low for both species. In addition, infection intensity was associated with clam size (proxy of age), but not with clam densities. Previous studies have also failed to find an association between *P. chesapeaki* infection and mortality (Bushek et al. 2008). However, this study was unable to disentangle the impact of disease on mortality (which would manifest as a negative relationship between bivalve density and disease) from density-dependent disease dynamics (which would manifest as a positive relationship between bivalve density and disease). To our knowledge, this study documents the most complete record of *Perkinsus* sp. infection in *M. arenaria* and *T. plebeius* to date, but impacts of disease mortalities on depleted clam populations of Chesapeake Bay remain to be carefully evaluated by rigorous experimental investigations of disease pathogenesis and outcomes.

Several major storms occurred during the study period, including Hurricane Sandy, which made landfall in New Jersey on October 29, 2012 and caused major flooding throughout Chesapeake Bay (Kunz et al. 2013), and Hurricane Lee in 2011, which was a major inflow event (Hirsch 2012). In contrast to Tropical Storm Agnes, these storms produced no noticeable mortality in our survey of *M. arenaria* and *T. plebeius* in the Chesapeake Bay. Although some of these storms caused extensive loss of human life and damage to infrastructure (Smith & Katz 2013), benthic ecosystems may be largely resistant to such disturbances. However, extreme storms have caused mass mortality of bivalves (Lochead et al. 2012, Freeman et al. 2013), and it is possible that these storms did not produce environmental conditions severe enough to produce a response in the bivalves surveyed as part of this study.

The lack of information on *T. plebeius* long-term abundances is an impediment to understanding the decline of this species and the consequences of this decline. Without landings records or fisheries-independent indices of *T. plebeius* abundance, it is difficult to assess the health of the population. In addition, basic population biology information for *T. plebeius* populations in the coastal U.S. is often lacking. Investigations of some populations of *T. plebeius* in Argentina (Lomovasky et al. 2016) and Brazil (da Silva et al. 2015) have established valuable baselines for the species. The ecological and commercial importance of this species, as well as its role harboring *Perkinsus* sp., warrant future studies on growth, reproduction, tolerance to environmental conditions, disease-related mortality, and distribution within U.S. waters.

### Conclusions

It is likely that habitat loss, low recruitment, and predators are major factors keeping *Mya arenaria* and *Tagelus plebeius* at low densities in Chesapeake Bay, though continued harvest, which was not addressed in this study, may play a role as well. We found no evidence suggesting densities of *M. arenaria* or *T. plebeius* are negatively impacted by *Perkinsus* sp. infection prevalence or intensity, though the direct effect of *Perkinsus* sp. infection on bivalve mortality remains to be addressed. Extremely low densities of *M. arenaria,* decimated after tropical storm Agnes in 1972 (Cory & Redding 1976, Haven et al. 1976), and a susceptibility of this thin-shelled species to predation by blue crabs, likely fuel a feedback loop that leads to high per-capita rates of predation, which works to keep *M. arenaria* populations at low levels. In recent years, similar dynamics may have been at work in the upper Chesapeake Bay, where populations of *T. plebeius* have reached low densities through fishing pressure or possibly disease. Since *M. arenaria* and *T. plebeius* are preferred prey items for major predators such as *Callinectes sapidus* and *Rhinoptera bonasus* (Blundon and Kennedy, 1982b; Fisher, 2010), it is unlikely that predator switching will provide much relief. However, both species appear able to take advantage of refuge provided by complex habitats. Habitats such as seagrass and shell hash allow both bivalve species to persist at a low-density refuge, which may be stable. Further work should focus on predator-prey interactions, elucidating the existence and stability of low-density refuge habitats, the effect of population declines on ecosystem services such as filtration, and the likelihood that thin-shelled bivalve species may persist, recover, and sustain commercial fisheries in Chesapeake Bay.

## ACKNOWLEDGMENTS

We gratefully acknowledge the assistance given by the students and staff of the Community Ecology and Marine Conservation Biology labs at the Virginia Institute of Marine Science, many Maryland Department of Natural Resources staff members who collected and analyzed clams for abundances and diseases during 2000-2009, and the Fish and Invertebrate Ecology Lab at the Smithsonian Environmental Research Center. This material is based upon work supported by the National Oceanic and Atmospheric Administration grant numbers NA11NMF4570218 and NA07NMF4570326. C. Glaspie was supported by an Environmental Protection Agency (EPA) Science to Achieve Results (STAR) Fellowship, grant number FP91767501; and the National Science Foundation Administration GK-12 program, grant number DGE-0840804. M. Ogburn was supported by a Postdoctoral Fellowship awarded by the Smithsonian Environmental Research Center. This work is based on a PhD dissertation of C. Glaspie, supervised by R. Seitz. This paper is contribution number XXXX from the Virginia Institute of Marine Science, College of William & Mary.

## SUPPLEMENTARY TABLES

**Supplemental Table 1.** Fish species list and total number caught at sites in the lower and upper Chesapeake Bay. In the lower Bay, a total of 432 20-m tows were conducted with a modified crab scrap. In the upper Bay, a total of 96 500-m tows were conducted with an otter trawl.

| Common Name | Scientific name | Number caught |  |
| --- | --- | --- | --- |
|  |  | Lower Bay | Upper Bay |
| White catfish | *Ameiurus catus* | 0 | 1 |
| Bay Anchovy | *Anchoa mitchelli* | many | many |
| American eel | *Anguilla rostrata* | 3 | 0 |
| Silver perch | *Bairdiella chrysoura* | 518 | 0 |
| Spadefish | *Chaetodipterus faber* | 10 | 0 |
| Spotted seatrout | *Cynoscion nebulosus* | 5 | 0 |
| Weakfish | *Cynoscion regalis* | 3 | 0 |
| Gizzard shad | *Dorosoma cepedianum* | 0 | 2 |
| Mummichog | *Fundulus heteroclitus* | 3 | 0 |
| Skilletfish | *Gobiesox strumosus* | 35 | 0 |
| Channel catfish | *Ictalurus punctatus* | 0 | 1 |
| Pinfish | *Lagodon rhomboides* | 71 | 0 |
| Spot | *Leiostomus xanthurus* | 380 | 213 |
| Rainwater killifish | *Lucania parva* | 12 | 0 |
| Atlantic croaker | *Micropogonias undulatus* | 381 | 0 |
| White perch | *Morone americana* | 0 | 2095 |
| Striped bass | *Morone saxatilis* | 19 | 84 |
| Oyster toadfish | *Opsanus tau* | 1 | 0 |
| Summer flounder | *Paralichthys dentatus* | 77 | 0 |
| Harvestfish | *Peprilus paru* | 1 | 0 |
| Yellow perch | *Perca flavescens* | 0 | 2 |
| Norther puffer | *Sphoeroides maculatus* | 6 | 0 |
| Tonguefish | *Symphurus plagiusa* | 12 | 0 |
| Hogchoker | *Trinectes maculatus* | 183 | 0 |
| Spotted hake | *Urophycis regia* | 5 | 0 |

## SUPPLEMENTARY FIGURES

**Supplementary Figure 1.**
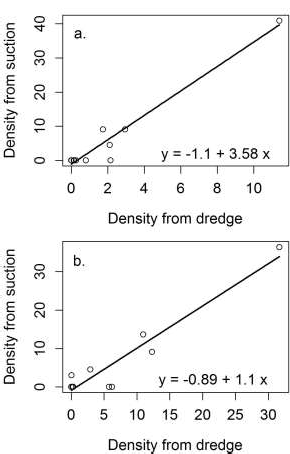
Relationship between density of (a) *Mya arenaria* and (b) *Tagelus plebeius* in suction samples and hydraulic dredge samples, including equations from simple linear regressions. The inclusion of the high-density data point in regressions did not significantly impact slopes for *M. arenaria* (z = 1.39, p = 0.17) or *T. plebeius* (z = 1.36, p = 0.18), as determined by a two-sided hypothesis test of z values calculated from model coefficients and standard errors (Clogg et al. 1995). High-density data points for both species do not come from the same location, and these densities were within the range of expected values for both species; thus, these data points were not omitted from analyses.

**Supplementary Figure 2.**
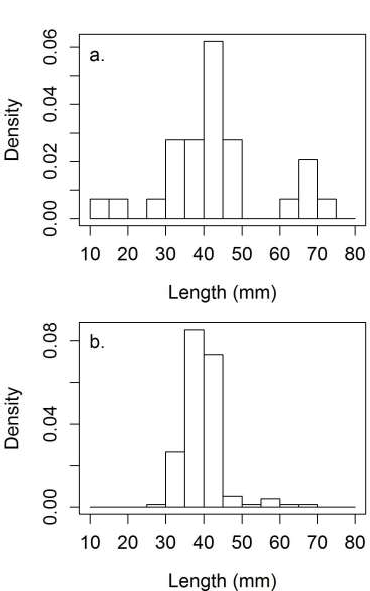
Size (length) density histograms for *Mya arenaria* captured in (a) suction samples and (b) hydraulic dredge samples.

**Supplementary Figure 3.**
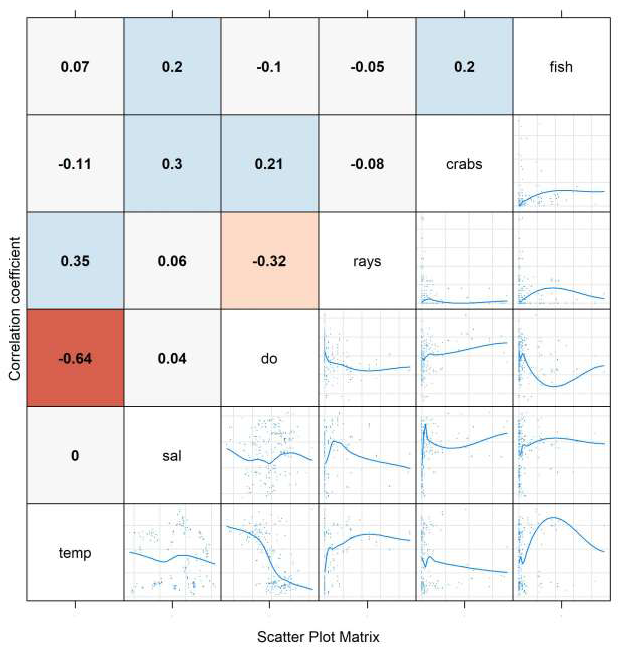
Pearson correlation coefficients (top panels) and scatter plots with splines (bottom plots) for all environmental and predator variables. Color denotes direction (negative = red, positive = blue) and magnitude (shading) of correlation coefficients.

**Supplemental Figure 4.**
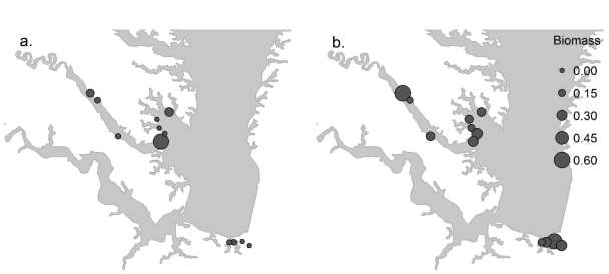
Lower Chesapeake Bay biomass (ash-free dry weight) of (a) *Mya arenaria* and (b) *Tagelus plebeius* captured in suction samples between fall 2011 and summer 2013. Point size is a linear function of mean biomass at each site: Radius = (4.0 × Density) + 1.

